# Fly wing evolution explained by a neutral model with mutational pleiotropy

**DOI:** 10.1101/2020.02.18.878595

**Authors:** Daohan Jiang, Jianzhi Zhang

**Affiliations:** Department of Ecology and Evolutionary Biology, University of Michigan, Ann Arbor, Michigan, USA

**Keywords:** mutational variance, phenotypic evolution, selection

## Abstract

To what extent the speed of mutational production of phenotypic variation determines the rate of long-term phenotypic evolution is a central question in evolutionary biology. In a recent study, Houle *et al.* addressed this question by studying the mutational variation, microevolution, and macroevolution of locations of vein intersections on fly wings, reporting very slow phenotypic evolution relative to the rates of mutational input, high phylogenetic signals of these traits, and a strong, linear correlation between the mutational variance of a trait and its rate of evolution. Houle *et al*. examined multiple models of phenotypic evolution but found none consistent with all these observations. Here we demonstrate that the purported linear correlation between mutational variance and evolutionary divergence is an artifact. More importantly, patterns of fly wing evolution are explainable by a simple model in which the wing traits are neutral or neutral within a range of phenotypic values but their evolutionary rates are reduced because most mutations affecting these traits are purged owing to their pleiotropic effects on other traits that are under stabilizing selection. We conclude that the evolutionary patterns of fly wing morphologies are explainable under the existing theoretical framework of phenotypic evolution.

## INTRODUCTION

A fundamental question in evolutionary biology is the extent to which the rate of long-term phenotypic evolution is determined by the rate of production of phenotypic variation by newly arising mutations (Lande 1976; Chakraborty and Nei 1982; Hill 1982; Lynch and Hill 1986; Lynch 1990; Schluter 1996; Wagner and Altenberg 1996; Futuyma 2010). This question likely has different answers for different traits. At one extreme are purely neutral traits whose evolutionary rates are dictated by the rates with which phenotypic variations originate via mutation. At another extreme are traits subject to strong positive selection such that their evolutionary rates are primarily determined by the strength, duration, and frequency of Darwinian selection instead of mutation. The lack of empirical answers to this question is in a large part owing to the scarcity of suitable data to address this question, because such data require the information of the same phenotypic traits from mutation accumulation lines, natural conspecifics, and different species.

Recently, Houle *et al.* addressed the above question by studying the evolution of locations of vein intersections on fly wings in the past 40 million years after inspecting over 50,000 wings from more than 100 Drosophilid species (Houle et al. 2017). They reported that (1) the rate of phenotypic evolution is orders of magnitude lower than the neutral expectation given the mutational variance, (2) the phylogenetic signals of most of these phenotypic traits are high, and (3) the evolutionary rate of a trait is linearly correlated with its mutational variance. Houle *et al*. examined nine existing models of phenotypic evolution but found none that is consistent with all of the above features. For instance, a neutral model of phenotypic evolution is consistent with a linear correlation between evolutionary rate and mutational variance and a high phylogenetic signal, but cannot explain the slow evolutionary divergence observed (Lynch and Hill 1986; Lynch 1991). Models consistent with a low evolutionary rate, however, predict nonlinear relationships between evolutionary rate and mutational variance and/or weak phylogenetic signals (Houle et al. 2017). After exhausting all existing models, Houle *et al*. suggested that their observations may be explained if “most mutations cause deleterious pleiotropic effects that render them irrelevant to adaptation, and, more importantly, the proportion of mutational variation that is deleterious is similar for all traits” (Houle et al. 2017). This proposal, however, is highly improbable because the chance that the proportion of deleterious mutational variation is similar for some 20 different traits is exceedingly low.

Here we demonstrate that the reported linear correlation between mutational variance and evolutionary divergence among the fly wing traits is an artifact resulting from the use of a biased method and that applying an unbiased method reveals a non-linear relationship. We further show that patterns of fly wing evolution are explainable by a simple model in which the focal traits are neutral but most mutations affecting these traits are purged due to their pleiotropic effects on other traits that are under stabilizing selection. Importantly, our model does not require the improbable assumption in Houle *et al.*’s proposal that all wing traits are equally impacted by the deleterious pleiotropic effect.

## MATERIALS AND METHODS

### Comparison of covariance matrices

To compare matrices representing mutational variances (*M*) and evolutionary divergence (*R*) of the same set of traits, Houle *et al*. first rescaled the matrices so that they have the same trace, which is the sum of diagonal elements. That is, they set 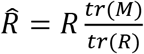, where *tr*(*M*) and *tr*(*R*) are traces of *M* and *R* matrices, respectively, and *Ȓ* is the rescaled *R* matrix. They then computed 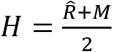. *M* and *Ȓ* were then converted to *K*^*T*^*MK* and *K*^*T*^*ȒK*, respectively, where *K* denotes the matrix comprising the eigenvectors of *H*. The diagonal elements of *K*^*T*^*MK* and *K*^*T*^*ȒK* were then compared to obtain a scaling exponent between *M* and *R*. Comparisons between *M* and *G* (the matrix summarizing phenotypic variances caused by additive effects of standing genetic variations) and between *R* and *G* were performed likewise.

Our new method is identical to Houle *et al*.’s method except that *K* is derived solely from *M* (when comparing *M* with *R* or *G*) or *G* (when comparing *G* with *R*) instead of *H*. The number of eigenvectors of *K* along which variances were compared is the same as the number of orthogonal traits plotted in the corresponding comparison of matrices in Houle *et al.* (2017). That is, regressions were performed using the first 18 eigenvectors of *K* when *M* was compared with *R* or *G* and the first 17 eigenvectors of *K* when *G* was compared with *R*.

### Examination of matrix comparison methods using simulation

To evaluate the performances of Houle *et al*.’s method and the new method in comparing *M* with *R* matrices, we simulated multivariate trait evolution on the phylogenetic tree of Drosophilid species used by Houle *et al*. and then analyzed the simulated data using the two methods, respectively. To ensure that the simulation is realistic and relevant, we used the *M* matrix estimated from the fly wing traits (Houle and Fierst 2013) as the true *M* matrix (denoted as *M*_true_ hereafter) in the simulation. Although the empirically estimated *M* matrix (Houle and Fierst 2013) may not be identical to the true *M* matrix, it is presumably similar to the true *M* matrix in terms of the structure. In each simulation, the matrix describing trait evolution, *R*_true_, was set to equal *wF* + (1 − *w*)*M*_true_, where *F* is a random matrix independent of *M*_true_ and *w* is its weight that ranges from 0 to 1. *F* was obtained by first generating a correlation matrix using the *rcorrmatrix* function of the R package *clusterGeneration* (Qiu and Joe 2015) and then converting it to a covariance matrix. Diagonal elements of *F* were sampled from pre-specified gamma distributions and then rescaled to have the same mean as the diagonal elements of *M*_true_. We sampled diagonal elements of *F* from gamma distributions with shape parameters (*k*) equal to 0.05 such that the skewness of the distribution of the diagonal elements of *R*_true_ is similar to that observed from the empirically estimated *R* matrix. Similar results were obtained when we used *k* = 0.025 or 0.1. The multivariate phenotype at each node was obtained by *X* = *X*_A_ + *l*∆*X*, where *X*_A_ is the phenotype at the node immediately ancestral to the focal node, *l* is the length of the branch connecting them, and ∆*X* is a vector sampled from the corresponding multivariate normal distribution of *R*_true_. For each combination of *w* and *k*, we performed 50 simulations, each with an independently simulated *F* as well as the corresponding *R*_true_.

After each simulation, we compared estimates of *M* and *R*, denoted as *M*_obs_ and *R*_obs_ respectively, using both Houle *et al*.’s method and the new method. *M*_obs_ was estimated from independent vectors taken from the distribution of *M*_true_. The sample size was set to be 150, because the empirical *M* matrix for the fly wing traits was estimated from 150 sublines (Houle and Fierst 2013). *R*_obs_ was estimated from the evolved phenotypes using the *ratematrix* function of the *geiger* package in R (Revell et al. 2007; Pennell et al. 2014). In each simulation, we also compared *R*_true_ with *M*_true_ at a set of orthogonal directions corresponding to the eigenvectors of *M*_true_ and considered the regression slope of log_10_(*R*_true_ variance) on log_10_(*M*_true_ variance) the true scaling exponent between evolutionary divergence and mutational variance.

### Simulation of evolution of neutral traits with mutational pleiotropy

#### Mutational input

For a neutral focal trait, mutations affecting the trait were generated per unit of time by simulation. The number of mutations followed a Poisson distribution with the mean equal to *λ*_M_, which is a random variable drawn from a gamma distribution with the shape parameter *k* = 0.5 and the scale parameter *θ* = 400. We set *k* = 0.5 because such a distribution is similar to some empirically observed distributions for mutationally independent orthogonal traits such as yeast cell morphologies (Ho et al. 2017) and fly wing morphologies (Houle and Fierst 2013). The phenotypic effect of a mutation on a trait followed a normal distribution with a mean of 0 and a standard deviation of 0.01. Therefore, *V*_*M*_, the expected phenotypic variance of the focal trait introduced by new mutations per unit time, equals *λ*_M_ x 0.01^2^. Before comparing mutational input and evolutionary divergence, we estimated *V*_*M*_ (*M* variance) for each trait from 150 independent samples taken from a normal distribution with the variance equal to the corresponding true value. These estimated *M* variances were compared with *R* variances.

#### Pleiotropic effects of mutations

We set the number (*n*) of traits genetically correlated with a focal trait to be the largest integer smaller than 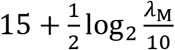. We assumed that *n* is a linear function of log_2_*λ*_*M*_ to impose a diminishing impact of *λ*_M_ on *n*, because when the focal trait is affected by multiple genes, it is unlikely that every one of them impacts a distinct set of additional traits. The probability that a mutation has an effect on a particular correlated trait was set to be 0.5. When the mutation was decided to have an effect, the effect size followed a normal distribution with a mean of 0 and a standard deviation of 0.01.

#### Fitness function and selection

Because the focal traits examined in the simulation were orthogonal, each simulation run considered only one focal trait and its genetically correlated non-focal traits. Upon the occurrence of a mutation on a background genotype, the fitness of the mutant was set to be 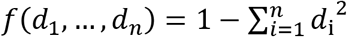, where *d*_i_ denotes the phenotypic distance from the optimum for the *i*th trait, *n* is the total number of traits correlated with the focal trait, and 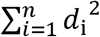 is the square of the Euclidean distance of the mutant from the multivariate optimum. If the fitness of the mutant relative to the fitness of the background genotype was lower than 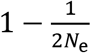, the mutation was removed by selection; otherwise, the mutation was accepted. Here, the effective population size *N*_e_ was assumed to be 10^6^ to model *Drosophila melanogaster* (Charlesworth 2009). When the Euclidean distance occasionally exceeded 1, we treated it as 1. At the end of each time unit, effects of all accepted mutations were added to the initial population mean phenotype. While the parameters used in the simulation presented are specific, we note that they are not unique in yielding results resembling patterns of fly wing evolution.

#### Phenotypic divergence and phylogenetic signal

For each trait with given *V*_M_ and *n*, we independently simulated its phenotypic evolution 50 times, all starting from the phenotypic optimum. Each simulation lasted for *t* = 2,000 units of time, after which the variance among the 50 replicates (*R* variance) was calculated at each time unit. Pearson’s correlation coefficient between time and *R* variance at the time was calculated to represent the phylogenetic signal. We note that the length of the simulation (*t*) has a negligible effect on the simulation results, because the focal trait is neutral and the *R* variance increases with time at a constant rate, as indicated by the high phylogenetic signal observed.

## RESULTS

### Houle *et al*.’s method of matrix comparison is biased

To investigate the scaling relationship between mutational variance and evolutionary divergence, one should compare the covariance matrices respectively representing the mutational inputs (*M*) and evolutionary divergences (*R*) of various traits along a set of orthogonal directions in the phenotypic space. For this purpose, Houle *et al.* used an eigenvector matrix *K* of the mean of *M* and *R* to determine the orthogonal directions. This practice could create a bias towards directions shared by *M* and *R* and inflate the correlation between log_10_(*M* variance) and log_10_(*R* variance). In theory, this potential bias can be avoided if *K* is derived solely from *M* such that the set of orthogonal directions are mutationally independent. We refer to this modified method as the new method.

To examine the potential bias of Houle *et al.*’s method and to compare its performance with that of the new method, we simulated phenotypic evolution along the fly phylogenetic tree used by Houle and colleagues. To be realistic, we treated the *M* matrix estimated from the fly wing data (Houle and Fierst 2013) as the truth (*M*_true_) and used it in our simulation. In the simulation, multivariate trait evolution followed a Brownian motion model, and the matrix describing trait evolution, *R*_true_, was obtained by taking a weighted average of *M*_true_ and an independent matrix *F* that adjusts the scaling between mutational input and evolutionary divergence (see Materials and Methods). After each simulation, we compared the estimated *M* and *R* matrices, denoted as *M*_obs_ and *R*_obs_, respectively, using Houle *et al.*’s method as well as the new method. We also compared *M*_true_ and *R*_true_ along a set of orthogonal directions corresponding to the eigenvectors of *M*_true_ and considered the regression slope of log_10_(*R*_true_ variance) on log_10_(*M*_true_ variance) the true scaling exponent.

When *M*_obs_ and *R*_obs_ are compared using Houle *et al*.’s method, the slope in the linear regression between log_10_(*R* variance) and log_10_(*M* variance) is close to 1 regardless of the true slope (Fig. 1a) and Pearson’s correlation coefficient between log_10_(*R* variance) and log_10_(*M* variance) exceeds 0.6 even when the true correlation coefficient is 0 (Fig. 1b). Clearly, Houle *et al*.’s method is uninformative and tells little about the true scaling between evolutionary divergence and mutational rate. By contrast, the slope and correlation coefficient estimated using the new method are close to the corresponding true values (Fig. 1). In theory, the new method may underestimate the slope because the *M* matrix used to obtain the orthogonal directions is *M*_obs_, which differs from *M*_true_ due to sampling error. Nevertheless, under the current simulation parameters, which are based on the actual fly wing data, the sampling error is sufficiently small to render the slope estimated by the new method reliable.

**Figure 1.**
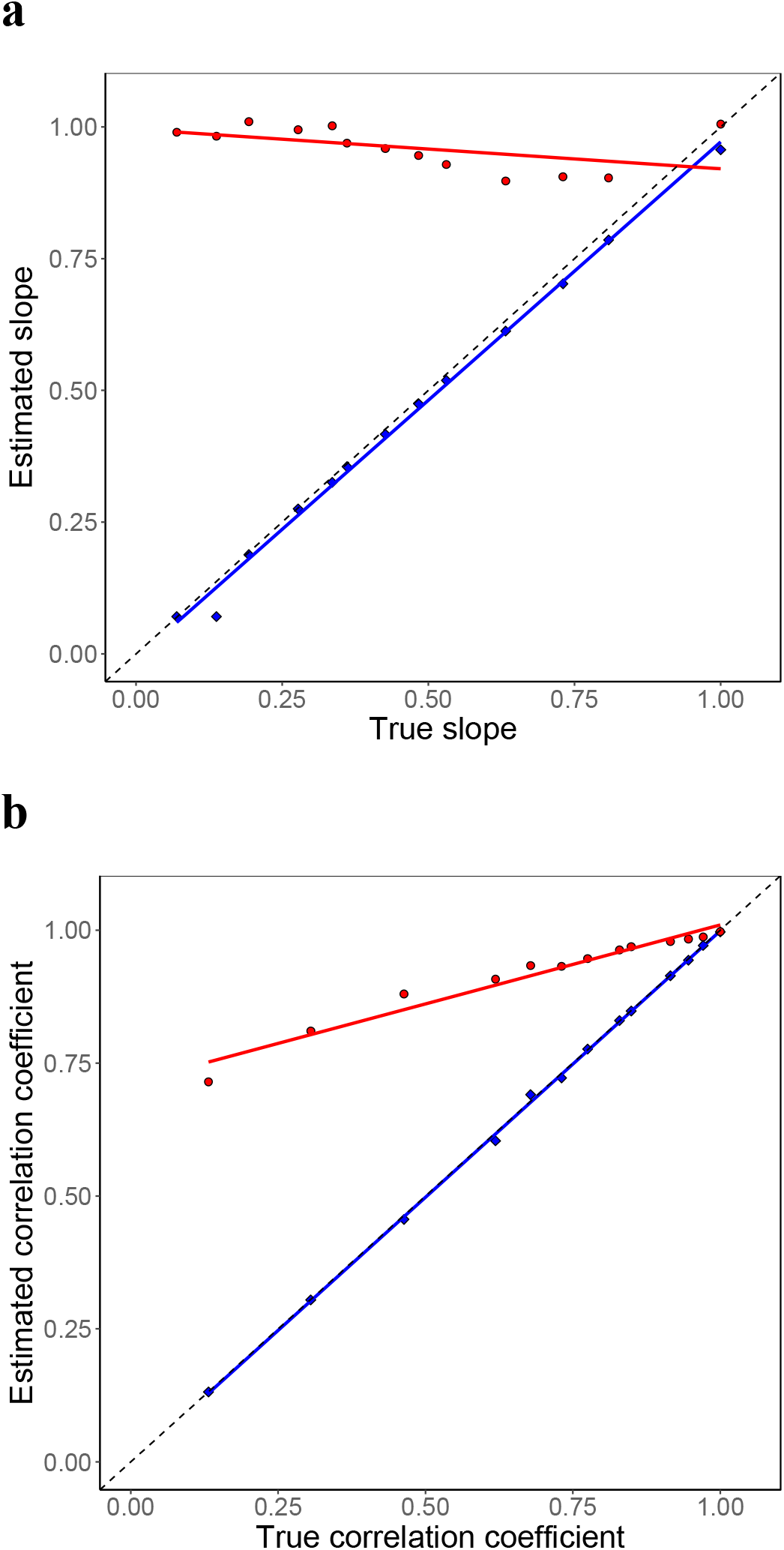
Slope of the linear regression and correlation between log_10_(*M*_obs_ variance) and log_10_(*R*_obs_ variance) of simulated data, estimated by Houle *et al*.’s method (red) and the new method (blue). **(a)** Slope of the linear regression between log_10_(*M*_*obs*_ variance) and log_10_(*R*__obs__ variance). **(b)** Pearson’s correlation coefficient between log_10_(*M*_*obs*_ variance) and log_10_(*R*_*obs*_ variance). Each dot represents the average slope or average correlation coefficient from 50 simulations with the same *w* parameter (see Materials and Methods). Colored lines in each panel are regression lines. The dashed line in each panel indicates the situation when estimates equal true values.

### Unequal constraints on mutationally independent traits

Using Houle *et al*.’s method, we reproduced their result of a slope of nearly 1 in the linear regression between log_10_(*M* variance) and log_10_(*R* variance) for fly wing traits (Fig. 2a). But our simulation suggested that this estimation is unlikely to be reliable. Indeed, when the fly wing data are reanalyzed using the new method, the slope reduced to 0.54, which is significantly smaller than 1 (*P* < 0.05, *t*-test; Fig. 2b). Applying the new method also caused a similar reduction in the slope of the linear regression between log_10_(*M* variance) and log_10_(*G* variance) (Fig. 2c-d), where the *G* matrix represents intraspecific phenotypic variations. The finding of a slope of approximately 0.5 for the linear regression between log_10_(*M* variance) and log_10_(*R* variance) or log_10_(*G* variance) indicates that *R* or G variance does not scale linearly with *M* variance. Rather, the positive impact of *M* variance on *R* or *G* variance gradually diminishes as *M* variance rises.

**Figure 2.**
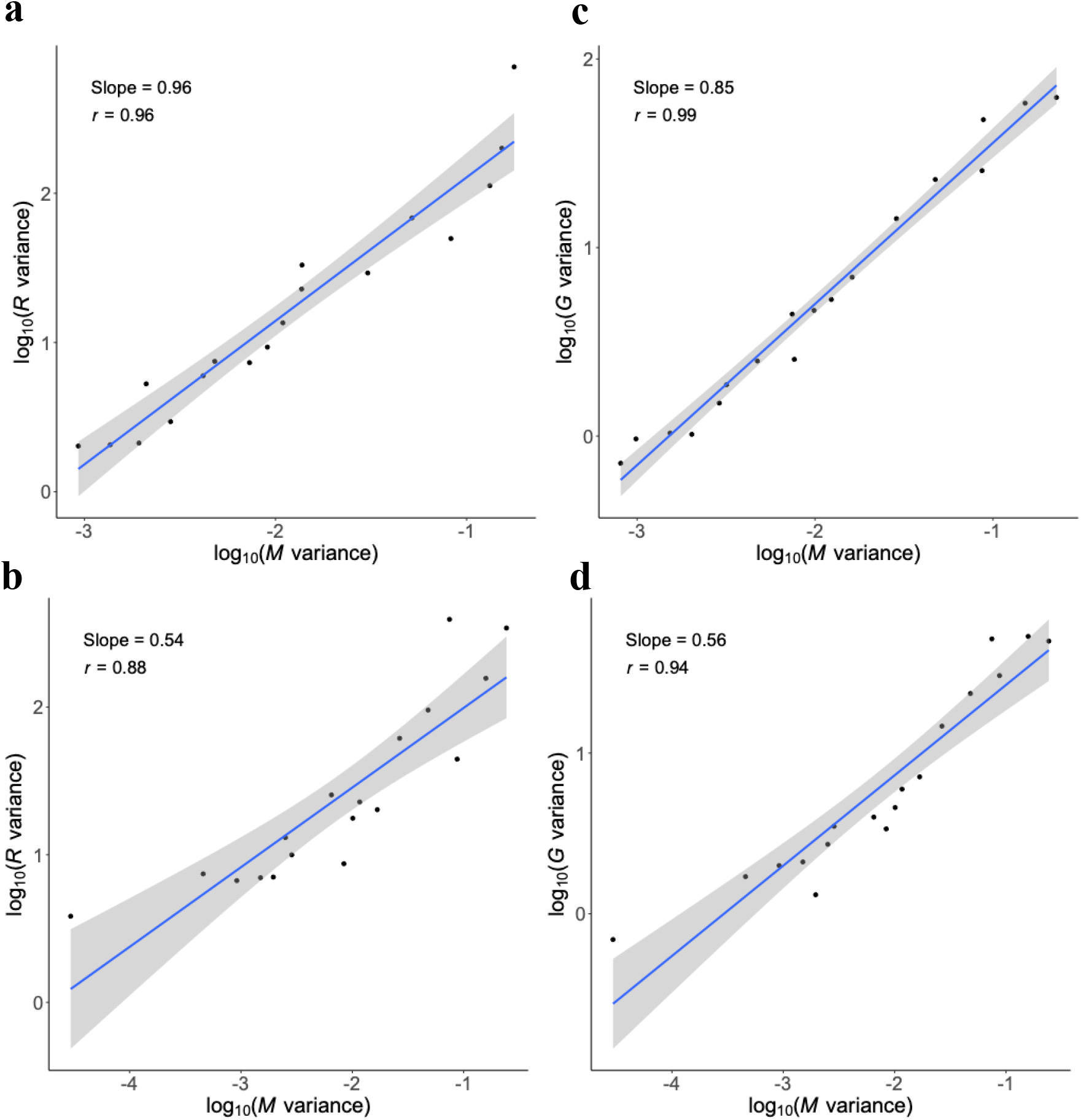
Linear regression between log_10_(*M* variance) and log_10_(*R* variance) or log_10_(*G* variance) along orthogonal directions for fly wing traits. (**a-b**) Linear regression between log_10_(*M* variance) and log_10_(*R* variance) estimated using Houle *et al.*’s method (**a**) or the new method (**b**). (**c-d**) Linear regression between log_10_(*M* variance) and log_10_(*G* variance) estimated using Houle *et al.*’s method (**c**) or the new method (**d**). In each panel, *r* stands for Pearson’s correlation coefficient, and the shaded region shows the 95% confidence interval of the regression. The number of orthogonal traits presented in each panel is the same as in Houle *et al*. (2017).

Interestingly, *G* and *R* are indeed similar in the structure. When the *G* and *R* variances are compared along the eigenvectors of *G*, the regression slope between log_10_(*G* variance) and log_10_(*R* variance) is 0.95, which is not significantly different from 1 (*P* = 0.48). This observation is consistent with the view that standing genetic variation has a profound impact on long-term evolution (Schluter 1996).

### A neutral model with mutational pleiotropy explains patterns of fly wing evolution

Houle *et al.* could not find a plausible model to explain fly wing evolution (Houle et al. 2017). Importantly, because the slope of the regression between log_10_(*M* variance) and log_10_(*R* variance) significantly deviates from 1, their proposal that all wing traits concerned are affected by pleiotropy to the same extent is not only implausible but also inconsistent with the data. Below we show that a neutral model with mutational pleiotropy can almost perfectly explain the above scaling between *M* variance and *R* or *G* variance as well as the first two observations of Houle *et al.* mentioned in Introduction. In our model, the focal wing traits are neutral, but mutations affecting the focal traits also influence other (unconsidered) traits that are subject to stabilizing selection (Turelli 1985; McGuigan et al. 2011). In addition, focal traits with higher *M* variances are likely influenced by more genes, which will likely affect more other traits. Consequently, focal traits with higher *M* variances are expected to be genetically correlated with more traits and impacted by greater mutational pleiotropy. Such a positive relationship may also arise from the positive correlation between the pleiotropic level of a mutation and its effect size on individual traits (Wagner et al. 2008; Wang et al. 2010).

Our model makes three predictions that are respectively consistent with the three patterns of fly wing evolution. First, a focal trait is expected to evolve more slowly than predicted from the *M* variance, because most mutations affecting the focal trait are selectively removed due to their deleterious effects on correlated traits. Second, because the focal trait itself is neutral, its divergence is unbounded, resulting in a high phylogenetic signal. Finally, the positive correlation between *M* variance and pleiotropy means that the fraction of mutations that are acceptable declines with *M* variance, creating a slope that is lower than 1 for the linear regression between log_10_(*M* variance) and log_10_(*R* variance) or log_10_(*G* variance).

To illustrate the above model predictions on long-term phenotypic evolution, we simulated the evolution of the population mean values of 20 orthogonal, neutral focal traits, each genetically correlated with a set of non-focal traits that are under stabilizing selection. Mutations were randomly generated per unit time and were accepted only if their fitness disadvantages are smaller than 1/(2*N*_e_) according to a phenotype-fitness mapping, where *N*_e_ is the effective population size. The simulation lasted for 2000 time units and was repeated 50 times per trait to create 50 replicate lineages. For each trait, the phenotypic variance among the 50 lineages at each of the 2000 time units was used to represent the evolutionary divergence (*R* variance) at that time, and its correlation with time is a measure of the phylogenetic signal. Details of the simulation are provided in Materials and Methods. The simulation results showed that, for most traits, the amount of phenotypic divergence is about four orders of magnitude lower than predicted from the total mutational input (Fig. 3a). In addition, all traits exhibited phylogenetic signals exceeding 0.9 (Fig. 3b). The slope of the linear regression between log_10_(*M* variance) and log_10_(*R* variance) is 0.52, which is significantly lower than 1 (*P* < 10^−10^, *t*-test; Fig. 3a). These results closely matched those observed in fly wing evolution, quantitatively verifying the validity and suitability of our model.

**Figure 3.**
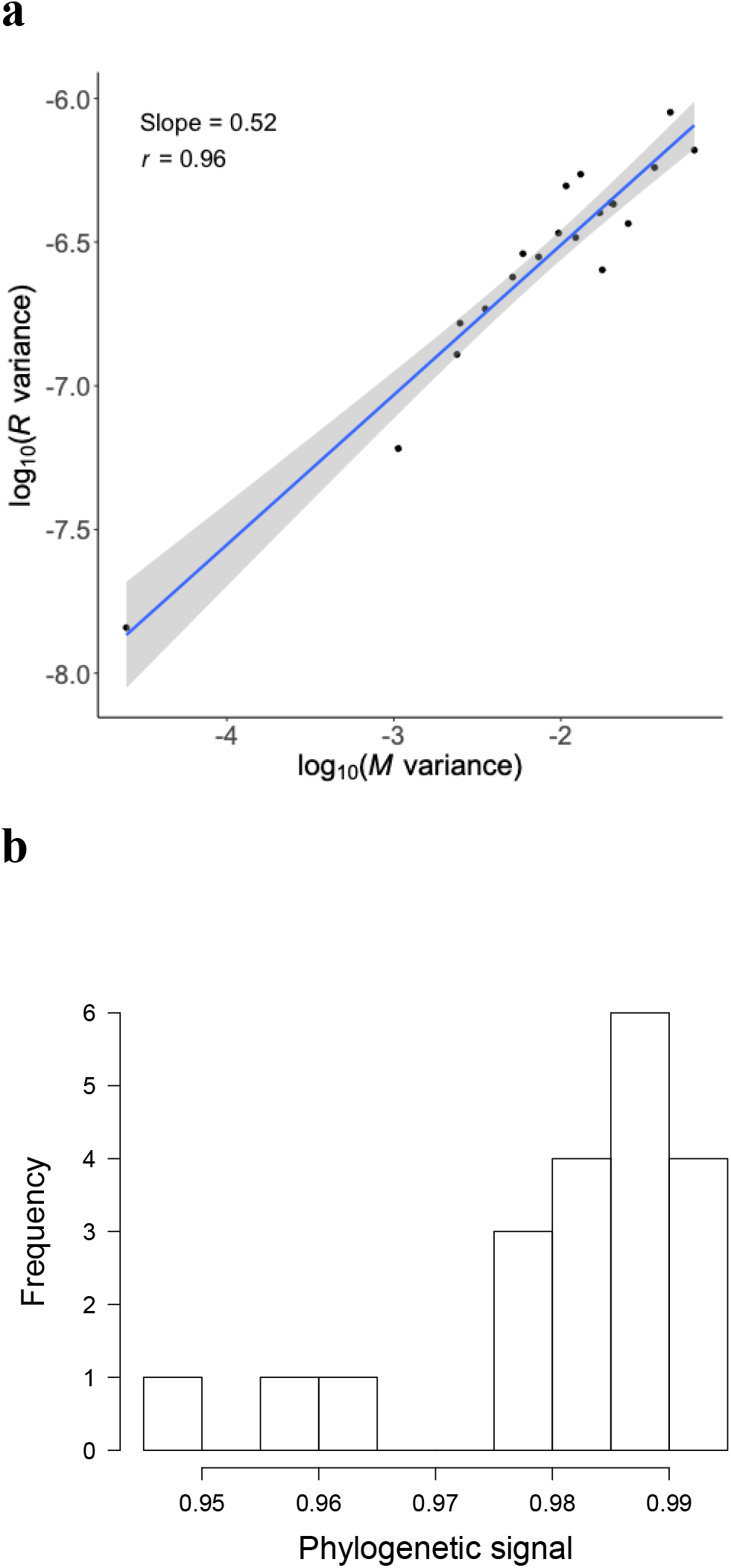
Patterns of phenotypic evolution observed from computer simulation of 20 orthogonal, neutral focal traits with mutational pleiotropy. **(a)** Linear regression between log_10_(*M* variance) and log_10_(*R* variance) upon evolution of 2000 time units. Presented are *M* and *R* variances per time unit. The shaded region shows the 95% confidence interval of the regression. **(b)** Distribution of the phylogenetic signals of the focal traits. The phylogenetic signal of a trait is measured by Pearson’s correlation between the evolutionary time and *R* variance at the time for the trait.

## DISCUSSION

In this study, we showed that the method used by Houle *et al.* to compare matrices is biased, resulting in the erroneous conclusion of a linear relationship between mutational variance and evolutionary divergence among fly wing morphologies. We demonstrated by computer simulation that a simple modification of their method yields virtually unbiased results under the parameters reflecting the fly wing data. Using the new method, we estimated that the scaling coefficient between mutational variance and evolutionary divergence is significantly smaller than 1, suggesting that the impact of the rate of mutational input on the rate of phenotypic evolution is not constant but declines with the rate of mutational input. That is, compared with traits with relatively low mutational inputs, those with relatively high mutational inputs do not evolve as rapidly as predicted linearly from their mutational inputs. With this finding, patterns of fly wing evolution are explainable by a model in which the wing traits are themselves neutral but mutations affecting the wing traits also affect other traits that are under various degrees of stabilizing selection. Our estimate of the scaling coefficient suggests that traits with higher mutational variances are subject to stronger mutational pleiotropy, and we offered potential mechanisms responsible for this relationship. Our evolutionary simulation under the above model is able to recapitulate all major patterns observed in fly wing evolution. Nevertheless, it is possible that the fly data also fit some other models. In particular, our results suggest the plausibility but do not prove that the fly wing traits are neutral. In fact, an expanded model in which the focal traits are neutral only within a range of phenotypic values can also explain fly wing evolution, provided that 40 million years of evolution under mutational pleiotropy has not reached the boundaries of this range. Regardless, our analysis suggests that fly wing evolution is explainable under the existing theoretical framework of phenotypic evolution.

The invaluable data collected by Houle *et al.* have allowed an unprecedented population genetic analysis of macroevolution of morphologies. To the best of our knowledge, no other large phenotypic data simultaneously comprising *M*, *G*, and *R* from long-term evolution exist. Only when many such data become available may we test the general applicability of our model or its expanded version in explaining phenotypic evolution, and only then can one tell whether the current theoretical framework of phenotypic evolution is generally correct.

## Acknowledgements

We thank Wei-Chin Ho and David Houle for valuable comments on earlier drafts. This work was supported in part by U.S. National Institutes of Health research grant GM103232 to J.Z.

